# Frequency-dependent effects of hip abductor vibration and surface translation on mediolateral sway

**DOI:** 10.1101/2025.11.17.688857

**Authors:** Sarah M. Rock, Maria C. Alvarez, Ethan B. Schonhaut, Jesse C. Dean

**Affiliations:** Department of Rehabilitation Sciences, Medical University of South Carolina, Charleston, SC, USA; Ralph H. Johnson VA Health Care System, Charleston, SC, USA

**Keywords:** Balance, Control, Posture, Proprioception, Somatosensation

## Abstract

The control of mediolateral standing balance is altered in many clinical populations, due in part to disrupted sensorimotor processing. Sensory perturbations, as evoked by musculotendon vibration, can provide insight into the contributions of individual sources of sensory feedback to this control. The purpose of this study was to investigate whether hip abductor vibration can elicit mediolateral sway at frequencies <1 Hz, which dominate standing posture. Secondarily, we quantified the effects of mediolateral surface translations in this frequency range. Participants (n=12) without neurological or orthopedic conditions completed a series of standing trials in which we quantified center of pressure motion. In a subset of trials, time-varying hip abductor vibration was delivered, with vibration intensity following sum-of-sine trajectories with frequency content from 0.1-0.9 Hz. In other trials, the standing surface translated mediolaterally following similar trajectories. Participants were not provided instructions regarding how to respond to the stimuli. Vibration significantly increased mediolateral sway for frequencies greater than 0.5 Hz, while surface translation caused substantial sway increases throughout the investigated frequency range. The effects of vibration were observed in the frequency range typically interpreted as reflecting feedback-driven corrective responses, suggesting the potential of this approach to influence balance performance through sensory augmentation.

## Introduction

The sensorimotor control of mediolateral standing balance is often altered in individuals with mobility deficits. Compared to neurologically-intact control participants, the magnitude of mediolateral sway during standing is typically increased among clinical populations with balance issues, including individuals with subacute and chronic stroke,^1,2^ diabetic peripheral neuropathy,^3^ cerebral palsy,^4^ and Parkinson’s disease.^5^ Greater mediolateral sway has also been linked with increased fall incidence among older adults,^6^ although with mixed predictive results in patients with neurological injuries.^7–9^ The discriminative ability of mediolateral sway to identify fallers can be increased when the control of balance is challenged, whether by closing the eyes,^6^ standing with a narrow stance width,^10^ or unpredictably translating the standing surface.^6^ In several cases, mediolateral sway has been found to be superior to anteroposterior sway in terms of predicting falls^6^ and differentiating between clinical population and control groups,^3–5^ justifying the need to identify the unique causes of altered frontal plane control.

One factor that can contribute to mediolateral balance deficits is disrupted sensorimotor processing. Neuromechanical models of varying levels of complexity indicate that the control of mediolateral standing balance is dependent on multisensory feedback with inherent physiological delays.^11,12^ Relative to anteroposterior balance, frontal plane control appears to be more dependent on proprioceptive feedback when standing with typical stance widths.^13,14^ Studies spanning numerous clinical populations have reported a link between reduced proprioception accuracy and increased mediolateral sway,^4,15–18^ supporting the importance of lower extremity proprioception. However, the strength of these reported relationships can vary across postural testing conditions (e.g., eyes open versus closed).

Beyond sway magnitude, sensory feedback also influences the frequency content of sway. During quiet standing in both control and clinical populations, the large majority of sway power occurs at frequencies below 1 Hz.^19–21^ Prior work has suggested that sway frequencies above ∼0.3-0.4 Hz reflect feedback-driven corrective responses, whereas lower sway frequencies are primarily due to slow movement of the body’s inertia,^22^ errors in the estimation of body state,^23^ or a changing reference posture.^24^ Changes in feedback-driven sway (frequencies>∼0.4 Hz) appear particularly important for balance; among people who have experienced a stroke, those who exhibited increased sway in this frequency range during quiet standing also tended to have poorer balance responses to perturbations.^25^ Most prior research in this area has focused on anteroposterior sway, rather than mediolateral, raising the question of whether similar frequency dependence is present in the frontal plane.

Compared to simply measuring postural sway during quiet standing, sensory perturbations allow more direct assessment of the extent to which specific sources of sensory feedback contribute to balance control. For example, the roles of visual and vestibular feedback in mediolateral balance have been assessed using moving visual surrounds^26,27^ and galvanic vestibular stimulation,^28,29^ respectively. Musculotendon vibration is a powerful stimulus to evoke proprioceptive feedback, and thus has often been used to investigate the role of this feedback source in the sensorimotor control of standing balance, as recently reviewed.^30^ Specific to mediolateral balance control, sustained vibration of the hip abductors causes sway away from the vibrated side,^31,32^ consistent with the importance of this muscle group in frontal plane posture^33^ and the evoked perception of lengthening of the vibrated muscle.^34,35^ Additionally, hip abductor vibration with time-varying intensity can increase mediolateral sway in a targeted frequency range of 0.45-0.65 Hz.^36^ While this result suggests that artificial proprioceptive feedback can shape the ongoing control of mediolateral sway, it is presently unclear whether such effects are present throughout the range of typically observed sway frequencies.

The primary purpose of this study was to investigate whether hip abductor vibration can elicit targeted mediolateral sway throughout the typical frequency range observed during standing posture (<1 Hz). We hypothesized that sinusoidal fluctuations in vibration intensity would cause increased mediolateral sway only at frequencies above 0.4 Hz, corresponding to the frequencies often cited as reflecting feedback-driven control.^22–24^ While our primary focus is on the effects of hip abductor vibration, we secondarily quantified the effects of continuous surface translations on mediolateral sway. Such translation methods are a common perturbation method to challenge standing balance,^37,38^ and the results will serve as an initial step toward our eventual goal of assessing the effects of hip abductor vibration on perturbed standing balance.

## Methods

### Participants

12 individuals (6F/6M; age=22±1 yrs; height=174±11 cm; mass=76±13 kg; mean±standard deviation) without neurological or orthopedic conditions participated in this study. The sample size was chosen based on the effect size (0.91) of mediolateral sway evoked by hip abductor vibration in prior work.^32^ With an alpha value of 0.05 for our primary paired comparisons between with and without vibration conditions, this sample size would achieve over 80% power. All participants provided informed consent using a form approved by the Medical University of South Carolina Institutional Review Board.

### Experimental Procedures

Participants completed a series of randomized-order 45-second experimental trials standing on a force plate capable of delivering prescribed surface translations (NeuroCom; Natus Medical Inc; Middleton, WI). For all trials, participants closed their eyes to increase their reliance on somatosensory feedback,^39^ crossed their arms to avoid contact with the vibrating tactors placed on the hips, and aligned their feet with a lightweight foam block in a position with the heels separated by 17 cm and the feet angled slightly laterally at 7°.^40^ Participants were instructed to stand still and relaxed, and were not given instruction on how to react to stimuli applied during trials. Participants completed 12 trials with hip abductor vibration, 12 trials in which the standing surface translated mediolaterally, and 3 control trials with neither vibration nor translation. Vibration or surface translation occurred during the middle 40-seconds of the trial, with the initial 2.5-second quiet period intended to ensure that participants were in a static posture before the stimuli began.

Vibration was delivered using tactors (EMS^2^; Engineering Acoustics Inc.; Casselberry, FL) secured over the bilateral hip abductors with silicone holders and elastic straps, directly superior to the greater trochanter and midway to the iliac crest.^41^ The vibration intensity varied over time, following sum-of-sines trajectories that were unpredictable to participants but consisted of well-defined frequency content (Figure 1). The frequencies of the sine waves ranged from 0.1 to 0.9 Hz, corresponding to the majority of mediolateral sway frequency content.^21^ Across trials, each sinusoidal frequency was delivered a total of four times. Increased vibration intensity took the form of higher vibration frequencies (up to 74 Hz) and displacements (up to 0.95 mm peak-to-peak).^42^ Increased vibration frequency in this range was used to elicit stronger responses from muscle spindles.^43^

**Figure 1.**
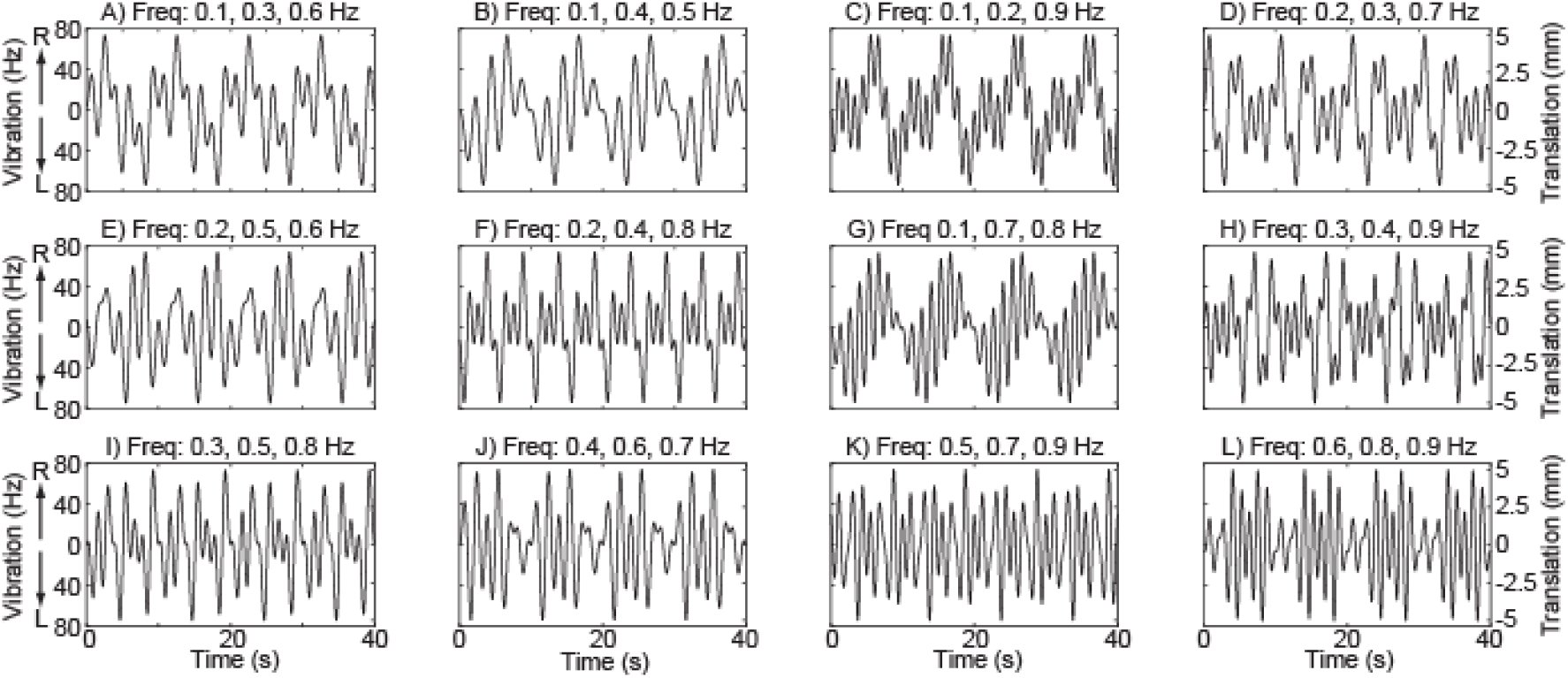
Time-varying trajectories of vibration intensity and surface translation. Trajectory profiles are identical for vibration (left y-axis) and translation (right y-axis) and consist of four repeating 10-second sum-of-sine patterns with the indicated sine wave frequencies and arbitrarily assigned phase shifts. Trajectories are ordered from lowest mean frequency to highest, with ties broken by geometric mean. For vibration intensity, positive values represent right side vibration while negative values represent left side vibration.

In translation trials, the standing surface underwent continuous mediolateral linear motion. The translation trajectories followed the same sum-of-sines pattern as the vibration intensity (Fig. 1), with displacement and velocity amplitudes sufficiently small (maximum peak-to-peak displacement = 10 mm; maximum absolute velocity = 25 mm/s) to not cause a loss of balance requiring a reactive response. As with the vibration trials, ensuring that the platform translation was unpredictable reduced the risk that participants would adapt their response.^44^ The vibration and translation trials were randomly interleaved, potentially increasing the likelihood that the sensory information evoked by vibration would be interpreted by participants as reflecting mediolateral sway.

### Data Collection and Processing

Our analyses focused on mediolateral motion of the center of pressure (CoP), as a CoP-based mechanism is believed to dominate the control of standing balance.^45^ We calculated CoP based on ground reaction force and moment data collected from the force plate, after compensating for inertial effects caused by platform translation.^46^ For each trial, we performed a Fast Fourier Transform (FFT) to quantify the frequency content of the mediolateral CoP location signal, focusing on frequencies below 1 Hz that dominate mediolateral sway.^21^ We also calculated the standard deviation of the mediolateral CoP displacement and velocity over the duration of each trial, common measures of sway performance.^21,47^

### Statistical Analyses

Our primary analysis investigated the effects of hip vibration on the frequency content of mediolateral sway, quantified as the FFT-derived sway amplitude at frequencies from 0.1-0.9 Hz, in 0.1 Hz increments. For each participant, we calculated the average sway amplitude at each of these frequencies for: 1) trials in which no vibration or translation occurred; 2) trials in which vibration was delivered, but the vibration trajectory did <u>not</u> include a component at the sway frequency of interest; and 3) trials in which vibration was delivered with a component at the sway frequency of interest. We used a repeated measures 2-way ANOVA to test whether sway amplitude was significantly affected by sway frequency (0.1-0.9 Hz) or vibration (3 conditions detailed above). In the event of a significant effect of vibration, we performed post-hoc paired t-tests at each of the frequencies of interest. Secondarily, we performed repeated measures 1-way ANOVAs to determine whether the standard deviation of mediolateral CoP displacement or velocity varied significantly across the trials without vibration or translation and the trials corresponding to the 12 vibration trajectories. Alpha values less than 0.05 were considered significant.

To explore the effects of surface translations on the frequency content of mediolateral sway, we repeated the statistical analyses described above but focused on trials with translation instead of vibration.

## Results

Time-varying hip abductor vibration influenced the frequency content of mediolateral sway. Sway frequency content for each vibration trajectory is illustrated in Figure 2. Increased sway amplitude with respect to the control condition was often observed at the frequencies targeted by the vibration intensity fluctuations. Across vibration trajectories, sway amplitude was significantly (p<0.001) influenced by sway frequency, with the most sway at the lowest frequencies (Figure 3A). Sway amplitude was also significantly (p<0.001) increased by the presence of vibration targeting that sway frequency (Figure 3A), in comparison to both control trials and trials with vibration that did not target that sway frequency. Post-hoc comparisons performed for individual sway frequencies found that compared to the control condition, vibration caused a significant (p<0.05) increase in sway amplitude at frequencies greater than 0.5 Hz.

**Figure 2.**
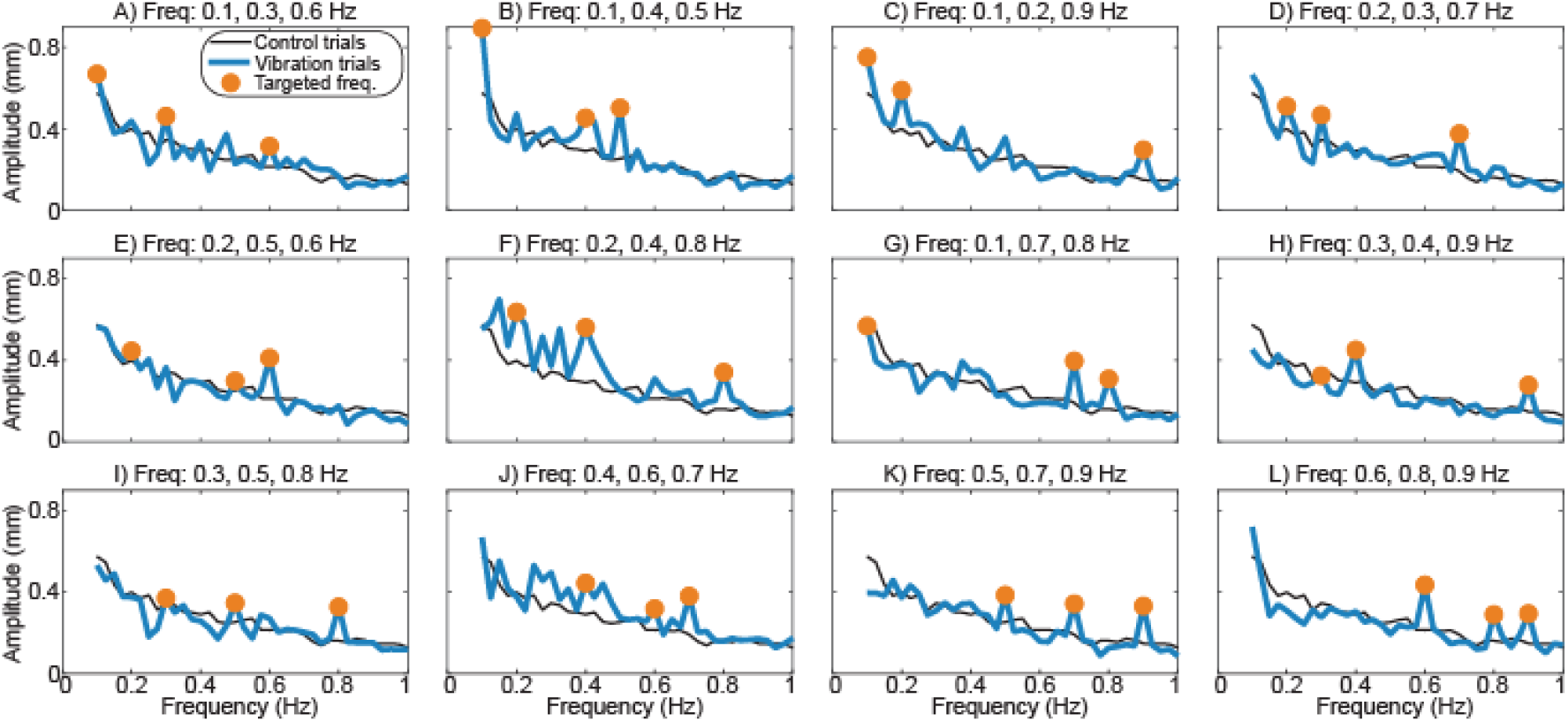
Effects of time-varying vibration on the frequency content of sway. The mean FFT-derived amplitudes of sway are plotted for frequencies from 0.1 to 1.0 Hz. The 12 panels illustrate sway behavior for the 12 vibration trajectories (thick lines), matching the order in Figure 1. For comparison, the mean sway behavior during control trials without vibration is also presented (thin lines). The three sway frequencies targeted for each trajectory are indicated by dots. Variability is not illustrated for visual clarity but is presented in Figure 3.

**Figure 3.**
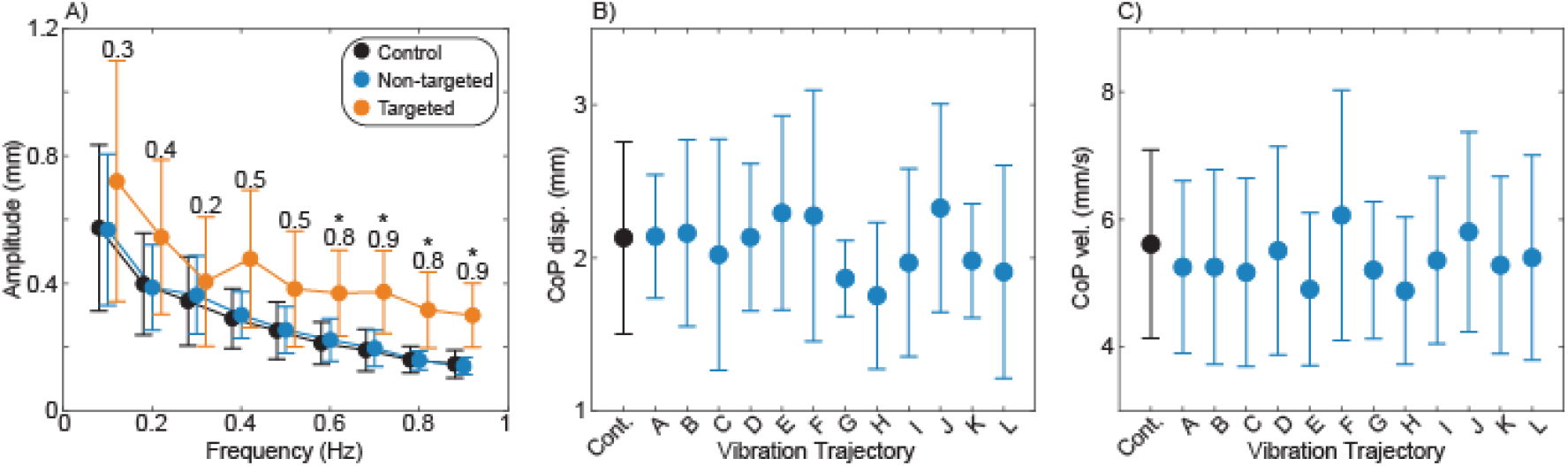
Group effects of vibration. A) Sway amplitudes were compared across frequencies ranging from 0.1 to 0.9. The sway amplitudes at each frequency are illustrated for three conditions: trials without vibration (Control); trials in which fluctuations in vibration intensity do not occur at this frequency (Non-targeted), and trials in which fluctuations in vibration intensity do occur at this frequency (Targeted). The Cohen’s d effect size for the Targeted condition compared to the Control condition is presented for each frequency. Asterisks (*) indicate a significant (p<0.05) difference between the Control and Targeted conditions. B) The standard deviation of mediolateral CoP displacement is presented for the Control condition and each of the twelve vibration trajectories. C) The standard deviation of mediolateral CoP velocity is presented for the Control condition and each of the twelve vibration trajectories. Across all panels, dots represent means and error bars represent 95% confidence intervals.

Despite the effects on sway frequency content, vibration had a minimal influence on traditional measures of mediolateral sway magnitude. No significant differences were observed across trials without vibration and with the 12 vibration trajectories in terms of either CoP displacement (p=0.61; Figure 3B) or CoP velocity (p=0.76; Figure 3C).

Surface translation had a major effect on mediolateral sway frequency content. Translation increased sway amplitude throughout the 0.1-0.9 Hz range, with particularly large increases at frequencies at which the translation occurred (Figure 4). Sway amplitude was significantly influenced by both sway frequency (p<0.001) and the presence of translation (p<0.001) (Figure 5A). Post-hoc tests found that across the range of individual sway frequencies, the presence of translation *at other frequencies* significantly increased sway amplitude (p<0.001). For sway frequencies above 0.1 Hz, further significant (p<0.001) increases in sway amplitude occurred when the translation included a component at the frequency under investigation.

**Figure 4.**
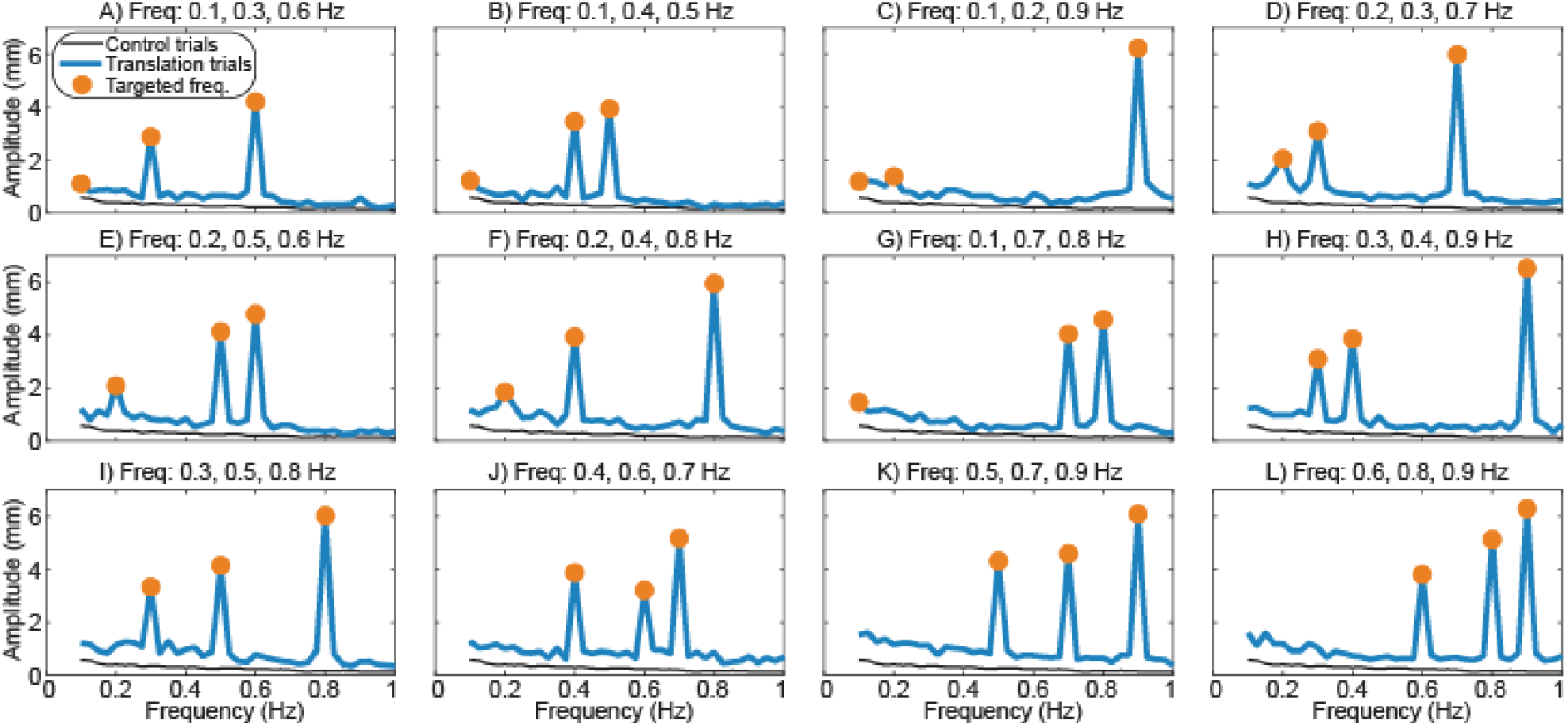
Effects of surface translation on the frequency content of sway. Figure structure matches Figure 2.

**Figure 5.**
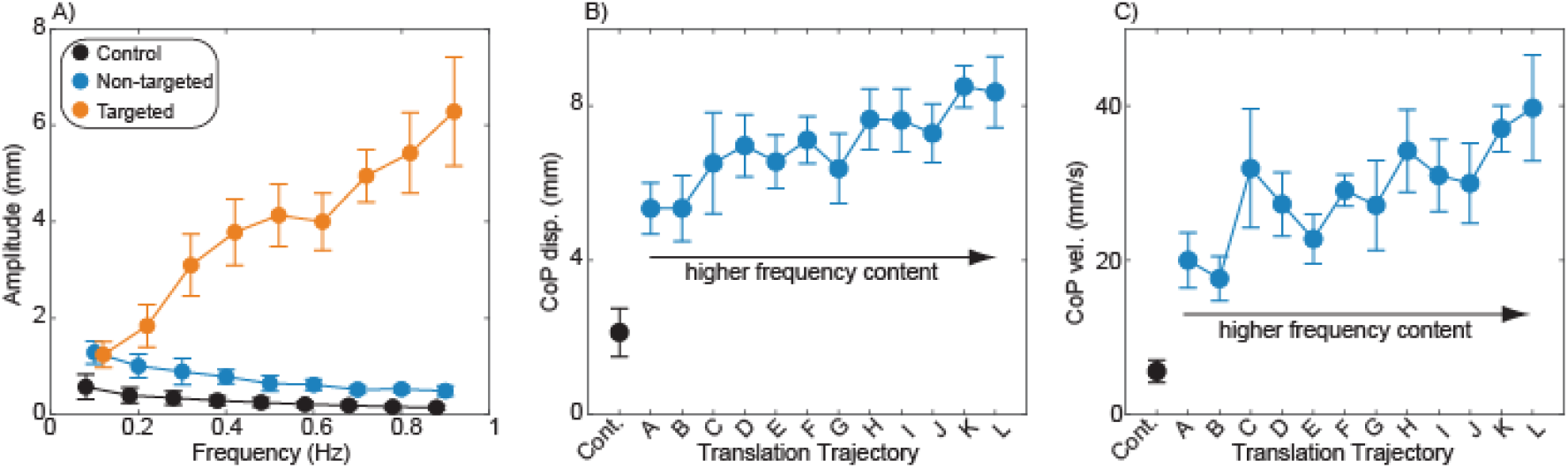
Group effects of surface translation. A) Paralleling Figure 3, sway amplitudes at each frequency are illustrated for trials without surface translation, with surface translations that do not occur at this frequency, and with surface translations that do occur at this frequency. The only comparison between conditions that did not achieve statistical significance was between Non-targeted and Targeted conditions at 0.1 Hz. All other comparisons had large effect sizes (Cohen’s d≥1.4), so are not illustrated here. B) CoP displacement standard deviation was significantly larger for all translations trajectories than for the control condition, and tended to increase for trajectories with higher frequency translation. C) CoP velocity standard deviation was significantly larger for all translations trajectories than for the control condition, and tended to increase for trajectories with higher frequency translation.

Surface translation also caused large increases in traditional measures of sway. Both mediolateral CoP displacement (p<0.001; Figure 5B) and CoP velocity (p<0.001; Figure 5C) varied significantly across trials with and without translation. Both CoP metrics were consistently higher for trials that included translation and tended to be greater for translation trajectories that included higher frequency components.

## Discussion

Both hip abductor vibration and surface translation altered the frequency content of mediolateral sway while standing. Our hypothesis was partially supported, as hip vibration caused larger increases in sway at higher frequencies, although statistical significance was only reached in the 0.6-0.9 Hz range. The sway evoked by surface translation was much larger than that caused by hip vibration, with clear increases in sway amplitude as translation frequency increased.

Artificial somatosensory feedback elicited increased mediolateral sway in a frequency range reflective of feedback-driven corrections.^22–24^ Specifically, we generally observed large (effect size [E.S.] ∼0.8) statistically significant effects in the 0.6-0.9 Hz frequency range, medium (E.S. ∼0.5) effects in the 0.4-0.5 Hz frequency range, and small (E.S. ∼0.3) effects with the slowest frequencies. Broader significant effects likely would have been detected with a larger sample size. This finding extends prior results^36^ to a wider frequency range, supporting further investigation of using hip abductor vibration to influence the feedback-driven control of mediolateral standing balance. While vibration increased sway amplitude at the targeted frequencies, it did not increase mediolateral CoP displacement or velocity, which are more traditional measures of sway. We attribute this lack of a global sway increase to the constrained effects of the delivered vibration; unlike with surface translation, hip vibration did not increase mediolateral sway at non-targeted frequencies, indicating a specificity of response that would be beneficial in a control paradigm.

The ability to elicit feedback-driven sway responses using hip abductor vibration suggests the potential of this approach to shape behavior through sensory augmentation. Sensory augmentation is broadly defined as the use of artificial sensory signals to provide useful information about body position or motion.^48^ Vibration has often been used in sensory augmentation systems that target balance,^49–51^ although typically in the form of tactile cues that users must attend to and cognitively process to generate a motor response.^48^ In contrast to most prior work, we did not instruct participants how to respond to the delivered vibration (e.g., when you feel vibration on your right hip, shift to the left). Instead, we relied on the natural physiological pathway through which abductor vibration is perceived as muscle lengthening that would accompany sideways sway.^31,32^ Extending the present open-loop control of hip vibration to a closed-loop system in which vibration intensity and location are controlled based on measures of real-time sway could have the potential to influence balance performance, as we have observed using a similar approach to shape foot placement during walking.^41,42^ Such an approach may be of particular value in populations with deficits in cognitive processing, as the increased cognitive demands needed to attend to cues can harm either cognitive or balance performance during dual tasks.^52–54^

Mediolateral CoP motion was highly sensitive to the frequency content of surface translations, with greater CoP motion accompanying higher frequencies. This clear relationship can be most simply attributed to the nature of CoP measurements, which reflect both the position and acceleration of the body’s center of mass.^33^ While the amplitude of the applied surface translation displacements was identical across frequencies, the translation accelerations increased with the square of the frequency. This larger effect of translation on CoP at higher frequencies is consistent with both mechanical expectations and prior empirical results.^55^ The relationship between surface translation and CoP motion is discussed in more detail in Appendix A, in which we explore the extent to which CoP trajectories can be predicted from surface translation patterns using simple mechanical models. As surface translations are commonly used to probe balance performance,^56–58^ the present results will help future studies to design mediolateral translation trajectories that elicit desired sway frequency characteristics.

Several important limitations of the present work must be addressed prior to considering clinical use. First, participants performed all trials with their eyes closed, which was intended to increase their reliance on somatosensory feedback that could be influenced by the vibration. It is thus possible that the effects of vibration would be smaller if participants had their eyes open. However, prior work found that hip vibration had a larger effect on mediolateral sway with eyes open than eyes closed, which may reflect increased muscular co-activation when the eyes are closed.^36^ Second, it is presently unclear whether hip abductor vibration would have similar effects on feedback-driven corrections among clinical populations with deficits in balance control. It is often assumed that members of such populations, such as those who have experienced a stroke, have an increased reliance on visual feedback^59^ and thus may not respond to proprioceptive stimuli. However, experimentally testing this assumption has revealed that the majority of individuals in this population were indeed responsive to hip abductor vibration delivered as a sensory perturbation.^60^ Finally, humans often reduce their responses to sensory stimulation over time when the stimulation does not align with other feedback sources and can be deemed unreliable.^61^ Therefore, the effects observed with the present open-loop control may actually be increased in a closed-loop context, where the delivered stimulation could be consistent with other sensory sources.

In conclusion, time-varying hip abductor vibration evokes mediolateral sway that is consistent with feedback-driven corrections used to ensure standing balance, even in the absence of instructions of how participants should respond to the stimuli. This result justifies future work extending this approach to closed-loop control, adding to previously investigated somatosensory augmentation methods that have primarily relied on cognitively interpreted tactile cues.

## Acknowledgements

This work was partially supported by a grant from the National Science Foundation (2242812) and by a grant from the National Institutes of Health (P30GM154630).

