## Appendix A for "Frequency-dependent effects of hip abductor vibration and surface translation on mediolateral sway"

**Introduction**

Balance control strategies during standing posture have long been assessed by applying mechanical perturbations and measuring participant responses.^1,2^ As presented in this manuscript’s main text, mediolateral translation of the standing surface has clear effects on the magnitude and frequency content of mediolateral sway, which allows basic quantification of perturbation responses. However, the relationship between an applied mechanical perturbation and the resultant sway has the potential to provide further insight into the underlying balance control strategy, a goal that has been facilitated by the use of mathematical models.^3^

While many prior models of standing balance have focused on anteroposterior motion, the control of mediolateral balance has also been investigated using models of widely varying complexity. At its simplest, the mechanical structure of the body during mediolateral sway has been analogized as a sliding mass,^4^ or a single-link inverted pendulum.^5–7^ More complex models have treated the legs, pelvis, and ground as a 4-bar linkage,^8^ or used a multisegment model that allows independent motion between the lower body (legs and pelvis) and upper body (trunk, head, and arms).^9^

In addition to variation in mechanical structure, prior models have also differed in terms of the types of feedback that contribute to model actuation. The forces or torques influencing body motion have been simplified as passive springs and/or dampers resisting body or joint displacement or velocity,^4,6,7^ and analogous to a proportional-derivative (PD) control loop. An alternative approach includes a constant time delay in position and velocity feedback,^8^ accounting for known physiological delays in sensorimotor control. Still more complex models combine passive mechanical properties with variable time-delayed feedback proportional to body motion and torques,^5^ or allow independent time delays for upper and lower body motion.^9^ Finally, simple models tend to include single parameters to represent the summed available feedback from all sensory sources, whereas others separate the estimated feedback into individual sources (e.g., vestibular, proprioceptive, tactile).^5,9^ Across model types, more complexity allows more detailed questions to be posed (e.g., regarding the specific contributions and timing of sensory feedback types), whereas simpler models can be assessed with less experimental data and fewer (albeit broader) assumptions.

The purpose of the exploratory analyses in this appendix was to identify the simplest model able to predict the major features of an individual’s center of pressure (CoP) motion from the applied surface translation trajectory. As we did not quantify individual joint or body segment kinematics, the mechanical structure of our models is necessarily simple, and we do not derive insight into the role of individual sensory feedback sources. Instead, our models investigate the effects of increasing complexity in terms of the types and timing of global feedback (i.e., stiffness and damping, delayed and non-delayed). We defined a successful model as one that predicted the majority of variation in CoP displacement and velocity trajectories in feed-forward simulations.

**Methods**

*Model Types.* We assessed the predictive ability of ten simple mechanical models, illustrated in Figure S1 in order of the number of fit parameters (see Model Fitting section below). All models treated the motion of the body’s center of mass (CoM) as a mediolaterally sliding mass^4^ with its motion driven by translation of the standing surface. We justified this simplification based on two considerations: 1) we did not measure body or joint kinematics, and would thus be unable to validate predictions of more realistic body motion; 2) with the small body motions elicited by the applied perturbations, the previously used 4-bar linkage model^8^ essentially simplifies to a sliding mass at a constant height if one assumes the body segments’ masses are concentrated at the CoM.

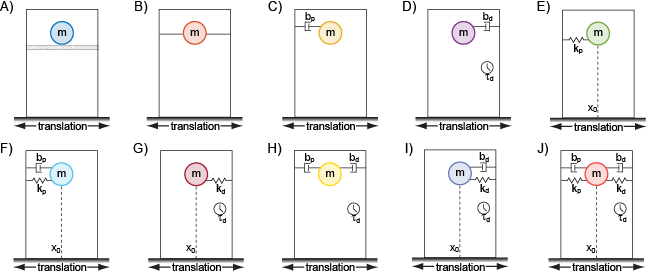

**Figure S1.** Models of increasing complexity were assessed, ranging from 0 fit parameters (A and B) up to 6 fit parameters (J). In panel A, the mass (m) simply slides on a frictionless surface. In panel B, the mass is held rigidly in place relative to the translating ground surface. In remaining panels, b_p_ represents passive damping, b_d_ represents delayed damping, t_d_ represents time delay, k_p_ represents passive stiffness, k_d_ represents delayed stiffness, and x_0_ represents zero point (position at which stiffness elements produce no force).

The simplest free sliding model (Fig. S1A) allows the CoM to translate freely as the surface moves under the body, with the CoM thus remaining fixed in space despite the applied perturbation. While seemingly unrealistic, this type of behavior has previously been qualitatively observed for anteroposterior sliding perturbations at relatively high frequencies.^10^ At the opposite extreme, the rigid body model (Fig. S1B) holds the CoM in a constant location relative to the moving support surface. The CoM motion thus matches the trajectory of the applied perturbation, similar to the previously observed behavior of “riding the platform” during low frequency anteroposterior sliding perturbations.^10^

For each of the remaining models, mediolateral forces are exerted on the CoM that are proportional to the relative displacement or velocity between the CoM and the support surface, corresponding to forces exerted in response to changes in body position or motion. This approach is consistent with the proposal that feedback-based balance responses are well predicted by CoM motion.^11^ Forces proportional to displacement are represented as springs, and can be thought of as proportional control. Models with springs also include a “zero point”, at which the force from the spring is zero. Forces proportional to velocity are represented as dampers, and can be thought of as derivative control. Several of these models (Fig. S1C, D, and F) include only damping and/or stiffness elements with no time delay, in which the generated forces are proportional to the CoM displacement or velocity at that instant in time. Following prior conventions,^5^ such parameters are considered “passive”, as such behavior could feasibly be achieved from passive mechanical properties of muscles or joints. Other models (Fig. S1D, G, and I) include only damping and/or stiffness elements with a time delay, in which the generated forces are proportional to CoM displacement or velocity at some earlier point in time. These models take into account potential sensorimotor delays. Finally, some models (Fig. S1H and J) include both passive and delayed elements.

*Model Fitting.* No model fitting was necessary for models A or B, as CoP trajectories were predicted directly from the surface translation trajectory. These models have thus zero fit parameters. The parameters of all other models were fit by relating mediolateral ground reaction forces (GRF) to estimated mediolateral CoM motion with respect to the standing surface. GRF were measured using the force plates participants stood on, with correction for inertial effects.^12^ Mediolateral CoM motion was estimated using a previously validated “zero-point-to-zero-point integration” method.^13^ Briefly, when the mediolateral GRF was zero, the CoP was defined to be directly under the CoM. Between these zero-crossings, mediolateral GRF was proportional to mediolateral CoM acceleration with respect to the stationary lab reference frame. Within these periods, we calculated mediolateral CoM velocity as the time integral of CoM acceleration, and mediolateral CoM displacement as the time integral of CoM velocity. Initial integration constants were calculated to ensure the displacement of the CoM over each time period matched the actual displacement of the CoP over this period. CoM velocity and displacement were then converted to the standing surface reference frame by subtracting motion of the support surface. GRF and CoM motion trajectories were upsampled to 1000 Hz for subsequent fitting.

Each model was fit to the GRF and CoM motion data for 10-second periods within trials while the standing surface was moving, as each of these trials included four repeated 10-second translation trajectories. Unlike some prior work,^14^ we did not average the experimental data across these repeated exposures before fitting. Such a strategy would reduce the risk of poorly-fit outliers (e.g., due to sway not caused by the surface translation), but also prevent the model from detecting changes in parameters over time. The best-fit parameters for each model were identified using a constrained linear least-squares approach. For example, Model C included only one parameter to be fit (passive damping; b_p_), solving the equation:

$$GRF\left( t \right)= {-b}_{p}\cdot\dot{x}_{COM}\left( t \right)$$

At the other end of the spectrum, the most complex model (Model J) included six fit parameters:

$$GRF\left( t \right)=-b_{p}\cdot\dot{x}_{COM}\left( t \right)-k_{p}\cdot\left( x_{COM}\left( t \right)-x_{0} \right)-b_{d}{\cdot\dot{x}}_{COM}\left( t-\tau_{d} \right)-k_{d}\cdot\left( x_{COM}\left( t-\tau_{d} \right)-x_{0} \right)$$

Here, *b_p_* is passive damping, *k_p_* is passive stiffness, *x_0_* is the zero point at which the springs exert no force, *b_d_* is the delayed damping, *k_d_* is the delayed stiffness, and *t_d_* is the time delay.

For models that included only passive damping and/or stiffness, the quality of the model fit was quantified as the R^2^ value between the predicted and actual GRF. For models with only delayed damping and/or stiffness, we solved the relevant equation for all potential t_d_ values ranging from 1 to 1000 ms. The optimal solution was chosen as that with the highest R^2^ value between the predicted and actual GRF. For models that included both passive and delayed parameters, possible t_d_ values were restricted to fall between 50 and 1000 ms to avoid excessive covariance between inputs. Damping and stiffness values were restricted to be greater or equal to zero, as negative values are unlikely to be physiologically realistic and consistently caused subsequent simulations to be unstable.

*Model Predictions.* For the free sliding model (Fig. S1A), the CoM remained in the same location in the lab reference frame, while the sliding surface translated beneath the body. The CoP trajectory on the force plate was thus simply predicted as the opposite of the surface translation trajectory. For the rigid body model (Fig. S1B), the CoM follows the same trajectory as the surface translation. Therefore, the applied translation acceleration is equivalent to the CoM mediolateral acceleration, and equal to mediolateral GRF divided by participant mass. The mediolateral location of the CoP (*x_CoP_*) was then calculated from the following equation:

$$x_{CoP}(t)=x_{COM}(t)-\frac{h}{g\cdot m}GRF(t)$$

Here, *h* is the height of the CoM (estimated to be 55% of participant height^15^) and *g* is the acceleration due to gravity. This equation assumes minimal angular motion of the model around the CoM, such that the CoP vector can be assumed to point through the CoM. CoP location is calculated as being under the CoM location, with an offset to account for CoM acceleration caused by the mediolateral GRF.

For all other models (Fig. S1C-J), we performed feed-forward model simulations to predict CoP trajectory based on the combination of support surface motion and model parameters identified as described in the prior section. Such feed-forward simulations in which future data values are not known are a more stringent test of model predictive accuracy than model fitting in which the entire data trajectory is included in the fitting process.^16^ For each 10-second period, these simulations were provided with the initial condition of CoM displacement and velocity from the estimated experimental data. We subsequently solved the delay differential equation for each model moving incrementally forward in time, using the dde23 function in Matlab (version 2024b) to predict CoM motion over the next 10 seconds. CoM trajectory values were then used to calculate GRF and CoP trajectories over this 10 second period, using the equation above.

*Assessing Model Prediction Accuracy.* For each 10-second period, the accuracy of model predictions was quantified by calculating the R^2^ value between the predicted and experimental CoP displacement traces. Similarly, we calculated R^2^ values between predicted and experimental CoP velocity traces, in which high-frequency motions are more apparent. We sought to identify the simplest model with average R^2^ values over 0.5 for both of these comparisons.

*Assessing Differences between Translation Trajectories.* Upon completion of the above analyses, we found that the model with delayed damping (Fig. 1D) was the simplest model that achieved our target criteria. We therefore focused our last set of analyses only on this model, testing whether damping and delay values varied across the 12 surface translation trajectories using repeated measures one-way ANOVAs.

**Results**

*Model Prediction Accuracy.* Prediction accuracy varied substantially across models for both CoP displacement and velocity (Fig. S2). The inclusion of a damping element had the most obvious effect, as all models without damping (A, B, E, G) made much poorer predictions of both CoP displacement and velocity. The simplest model that met our target criteria for prediction accuracy was the delayed damping model (Model D), with a median R^2^ value of 0.56 for both CoP displacement and velocity predictions. No appreciable increases in prediction accuracy were observed for more complex models. For comparison, we also present the R^2^ values for the fits (not feed-forward predictions) of CoM acceleration that were used to identify parameter values (Fig. S2). Here, the addition of model parameters always increased R^2^ values, although only slightly for the more complex models.

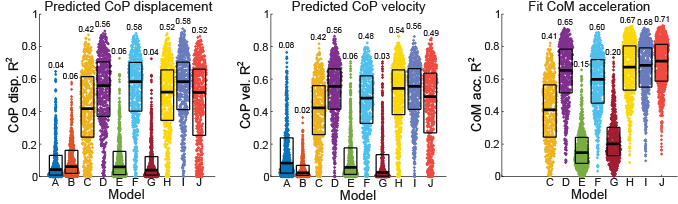

**Figure S2.** Coefficients of determination (R^2^ values) are illustrated for each model, participant, and 10-second period of sway as individual data points. Box plots indicate median, 25^th^ percentile, and 75^th^ percentile values, with median values included at the top of each data distribution.

For each model, the predicted CoP displacement and velocity trajectories are compared to the experimental trajectories for a representative participant in Figures S3 and S4. The predictions of the simplest models with no fit parameters have minimal resemblance to the experimental trajectories, primarily due to mis-timed CoP motion for the free sliding model (A) or excessive high-frequency content for the rigid model (B). Models that lacked a damping element (E, G) were similarly unable to predict the CoP motion; here, the model with delayed stiffness (G) was best fit with a minimal delay and thus was essentially identical to the model with passive stiffness (E). While the model with passive damping (C) predicted a CoP trajectory with a similar pattern to the experimental trajectory, it severely underestimated the peaks of motion. The primary features of CoP motion were predicted by the delayed damping model (D), with further increases in model complexity having only a minor effect on the predictions. The most notable weakness of model D was its inability to predict slow sway that was unrelated to the surface translation.

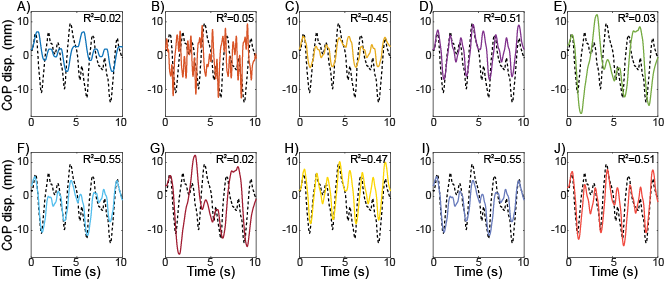

**Figure S3**. CoP displacement trajectories for a participant with R^2^ values close to the group median. Each panel illustrates the experimentally measured CoP displacement (dashed black line) and the feed-forward prediction from one of the mechanical models (solid colored line). Panel letters correspond to model type in Figure S1.

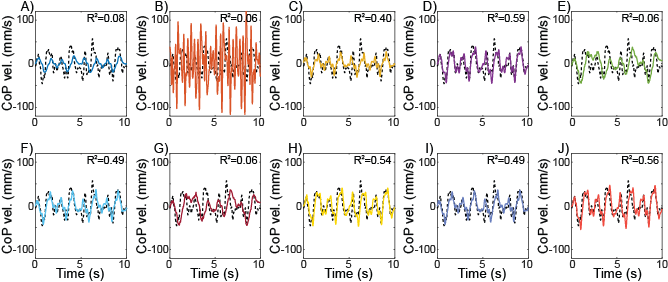

**Figure S4**. CoP velocity trajectories for a participant with R^2^ values close to the group median. Each panel illustrates the experimentally measured CoP velocity (dashed black line) and the feed-forward prediction from one of the mechanical models (solid colored line). Panel letters correspond to model type in Figure S1.

*Model Parameters.* Best-fit parameters for each mechanical model are illustrated in Figure S5. The parameter distributions for damping, stiffness, and delay were generally positively skewed, likely due to the set minimum value of zero. Overall, increasing model complexity (moving to the right on the plots) did not reduce parameter variability, providing an additional indication that more complex models do not necessarily provide clearer insight. The time delay parameter exhibited outliers on rare occasions for the delayed damping model (>500 ms for 1.4% of the 10-second periods). Such outliers became more prevalent with more complex models or when delayed stiffness was included.

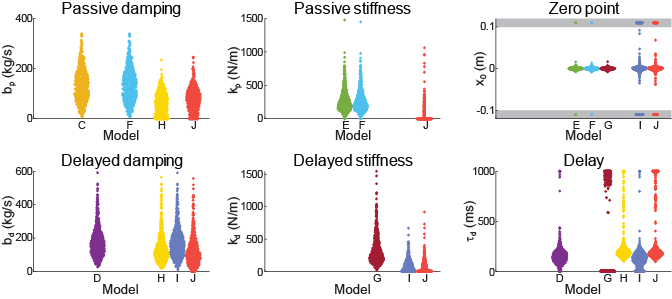

**Figure S5.** Best-fit parameter values for each model, participant, and 10-second period of sway are shown as individual data points. Extreme outliers for zero point values (magnitude > 0.1 m) are not presented on the same scale, and are included in the shaded areas.

*Translation Trajectory Effects.* The best-fit parameters of the delayed damping model (model D; the simplest model that met our target criteria) varied across translation trajectories. Damping magnitude varied significantly across trajectories (p<0.001), and tended to be smaller for trajectories with a higher root-mean-square (rms) frequency (Fig. S6). Similarly, delay varied significantly across trajectories (p<0.001), and was shorter for trajectories with a higher rms frequency (Fig. S6).

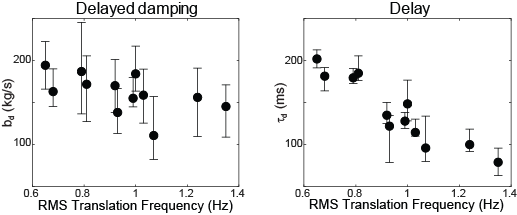

**Figure S6.** Both delayed damping magnitude and delay magnitude varied based on the translation trajectory. Data points represent median values for a given trajectory and error bars represent the 95% confidence interval of the median.

**Discussion**

A simple mechanical model including only two parameters (damping and delay) was able to predict the majority of the variation in mediolateral CoP displacement and velocity in response to surface translations. Added complexity in the form of passive damping, passive stiffness, or time-delayed stiffness did not appreciably improve prediction accuracy. The simplicity of this model prevents it from providing detailed neurophysiological insight into the control of standing balance. However, the model does allow potential insight into causal factors that could contribute to differences in CoP trajectories across populations (e.g., increased damping in older adults^17^), while requiring only data collected from a force plate.

Despite the relative simplicity of our delayed damping model, the primary results are consistent with more complex models. Adding passive (non-delayed) elements to our delayed damping model did not improve its predictions, corresponding to the prior finding that the effects of passive control parameters were much smaller than those of active (delayed) parameters.^8,9,17^ Beyond the presence of a time delay, the magnitude of the time delay identified by our simple model (median=147 ms; interquartile range=105-191 ms) was quite similar to the delays reported or assumed with more complex models (~114-220 ms).^5,8,9^ Finally, the observed variation in best-fit model parameters across translation trajectories also matches prior results. For example, an increased amplitude of frontal plane platform tilt has previously been found to cause increases in stiffness, but decreases in time delay.^5^ Such results could be attributed to either sensory reweighting or control nonlinearities, neither of which is a component of our simple model.

While our focus on simple models was intentional, this choice limited the extent to which our results can provide detailed insight into sensorimotor control. For example, our inclusion of only a single moving mass prevents predictions of the relative role of the lower extremities and trunk in body motion. However, prior work has found that lower body actuation had the largest effect on mediolateral sway for perturbation frequencies below 1 Hz (our perturbation range), whereas the trunk became more important at higher frequencies.^9^ Additionally, our use of a sliding mass rather than segments rotating around joints prevents us from basing feedback on joint angles, which may be a better representation of the feedback available from muscle spindles. Despite this limitation, prior work has found that feedback based on CoM motion produced quite similar results to more physiologically realistic joint feedback in terms of controlling frontal plane sway.^8^ Finally, while the proportional-derivative (PD) type of model used here is most common in studies of standing balance,^18^ alternative approaches such as those that involve optimal estimation theory^19^ explicitly include non-linearities and system noise that cannot be accounted for in our approach.
